# Three distinct profiles of visual category preference within the anterior temporal lobe

**DOI:** 10.64898/2026.03.11.711089

**Authors:** Chencheng Shen, Ben Deen

**Affiliations:** Department of Psychology and Brain Institute, Tulane University, New Orleans, LA 70118, USA

## Abstract

The primate visual hierarchy culminates in the anterior temporal lobe (ATL), yet the functional organization of this region remains poorly understood due to magnetic susceptibility artifacts that degrade fMRI signal quality. Here, we leverage a large-scale fMRI dataset from the Human Connectome Project (*N* = 830, male and female) to characterize visual category preferences within the ATL using a region-of-interest-based approach providing sensitivity in signal-compromised areas. Within the temporal pole (TP) and perirhinal cortex (PR), we identified three distinct profiles of visual category response, each replicated in an independent sample. TP showed a preference for objects over scenes, with a right-hemisphere bias toward animate objects (faces and bodies) over inanimate objects (tools). PR contained two functionally distinct subregions: one preferring faces over other categories, and one responding broadly to objects — particularly tools and bodies—over scenes. Analysis of spatial location revealed that, unlike the well-characterized topographic organization of category-sensitive regions in posterior occipitotemporal cortex, face- and object-preferring subregions of PR did not show a consistent spatial relationship, suggesting a patchy or “salt-and-pepper” organization. Resting-state functional connectivity analyses demonstrated that ATL subregions have dissociable connectivity profiles: TP showed preferential connectivity with transmodal default mode network regions, while PR subregions were more strongly coupled with occipitotemporal cortex, with each showing preferential connectivity with regions sharing similar category preferences. Together, these findings demonstrate that category sensitivity—a hallmark of the ventral visual stream—extends to the apex of the cortical visual hierarchy.

**Significance statement:** Understanding the functional organization of the visual system is a central goal of cognitive and systems neuroscience. The anterior temporal lobe (ATL) sits at the top of the brain’s visual processing hierarchy and is critical for recognizing objects and people, but its internal organization has remained poorly characterized due to technical limitations in brain imaging. Our findings resolve a key gap by demonstrating that the same organizational principle governing the well-studied posterior visual system—the segregation of responses by object category—extends all the way to the apex of the visual hierarchy. This lays essential groundwork for future research understanding interactions between visual and memory systems, and how high-level visual processing breaks down in neurological conditions affecting the ATL.

## Introduction

The primate cortical visual system is organized hierarchically: incoming visual information is first processed by early visual areas in the occipital lobe and then progresses anteriorly (Felleman & Van Essen, 1991). This visual hierarchy culminates in the hippocampal formation, preceded by surrounding structures in the anterior and medial temporal lobes. Extensive research on the functional organization of posterior, occipitotemporal parts of this system has revealed category-sensitivity as a dominant feature: different parts of cortex have preferences for particular ecologically relevant categories of visual input such as faces, bodies, manipulable objects or tools, and scenes (Downing et al., 2001; Epstein & Baker, 2019; Grill-Spector & Weiner, 2014; Kanwisher et al., 1997; Malach et al., 1995; Peelen & Downing, 2005). However, the organization of category-sensitive responses in the anterior temporal lobe (ATL) remains poorly understood, in part due to magnetic susceptibility artifacts that diminish fMRI signal quality from this part of cortex and make it difficult to study (Gorno-Tempini et al., 2002; Visser et al., 2010). To what extent does the ATL have a category-sensitive organization like posterior occipitotemporal cortex?

The ATL contains multiple regions that have visual responses and anatomical connections with the hippocampal formation. These include the temporal pole (TP, also termed area 38 or TG), positioned at the anterior tip of the temporal lobe, and perirhinal cortex (PR, also termed areas 35/36), positioned lateral to the collateral sulcus and anterior to the middle fusiform gyrus (Insausti et al., 1987; Moran et al., 1987; Suzuki & Amaral, 1994). Neuropsychological work has implicated both of these regions in aspects of visual perception and memory. TP dysfunction resulting from damage or neural degeneration can lead to deficits in recognizing objects and/or people from both visual and other sensory signals (Damasio et al., 1990; Gentileschi et al., 2001; Gorno-Tempini et al., 2004; Snowden et al., 2011). PR dysfunction can result in impaired memory for object identity (Buffalo et al., 1998) and has been argued to affect aspects of object perception (Barense et al., 2007; Murray & Richmond, 2001).

Despite anatomical and neuropsychological evidence for the role of the ATL in high-level vision, fMRI studies of category-sensitive visual responses have typically not reported significant responses in anterior regions (Grill-Spector & Malach, 2004; Kanwisher & Dilks, 2013). Nevertheless, some studies have suggested that category-sensitivity may extend to the ATL, including work finding responses to objects or faces in PR (Barense et al., 2010; Liang et al., 2013; Litman et al., 2009), and a face response in the ventral ATL (Axelrod & Yovel, 2013; Rajimehr et al., 2009; Tsao et al., 2008). In the macaque, where imaging of the ATL is not hampered by susceptibility artifacts, two face-preferring regions of ATL have been identified: one in TP and one in PR (Landi & Freiwald, 2017; Landi et al., 2021; She et al., 2024). Recent work has leveraged improvements in signal quality provided by multi-echo fMRI to identify potentially homologous face responses in human TP and PR, with the latter consistent in location with previously observed ventral ATL responses (Deen et al., 2024).

Here, we use a large-scale fMRI dataset (>800 participants) to explore whether the ATL contains category-preferring subregions, how they are anatomically organized, and how they interact with well-established areas in posterior occipitotemporal cortex. We leverage the large sample size and region-of-interest (ROI)-based analyses to increase sensitivity to detect responses in difficult-to-image brain areas. We find that the ATL contains three subregions with distinct category preferences: an object- or animate object-preferring region of TP, a face-preferring region of PR, and an object-preferring region of PR. The results suggest that the organizational principle of category sensitivity is not unique to mid-level visual areas, but rather extends to the apex of the cortical hierarchy.

## Materials and Methods

### Participants

This study used data from young adult participants from the HCP S1200 release, https://www.humanconnectome.org/study/hcp-young-adult/document/1200-subjects-data-release (Van Essen et al., 2013). The following exclusion criteria were applied: participants missing anatomical MRI, resting-state fMRI, or category localizer task fMRI data; participants with quality control issues identified by the HCP; participants with excessive head motion in functional data, defined as a mean framewise displacement across fMRI datasets of > .3mm. These exclusion criteria identified a subset of *N* = 830 participants from the full set of 1,206 participants.

In order to search for visual response profiles and replicate these profiles in an independent sample, we then split our participants into two subsets of *N* = 415, termed the *discovery* and *replication* groups. To ensure that groups were matched on proportions of males and females, and age distributions for each sex, we took participants with each age/sex combination and split them randomly across the two groups. The resulting age distributions are shown in Figure 1. Each sample included 227 females and 188 males, with age range 22 to 36 and mean age 28.6 years (Discovery) and 28.7 years (Replication). All participants were right-handed and had no history of medical condition. The study protocol was approved by the Washington University institutional review board, and participants provided written, informed consent.

**Figure 1:**
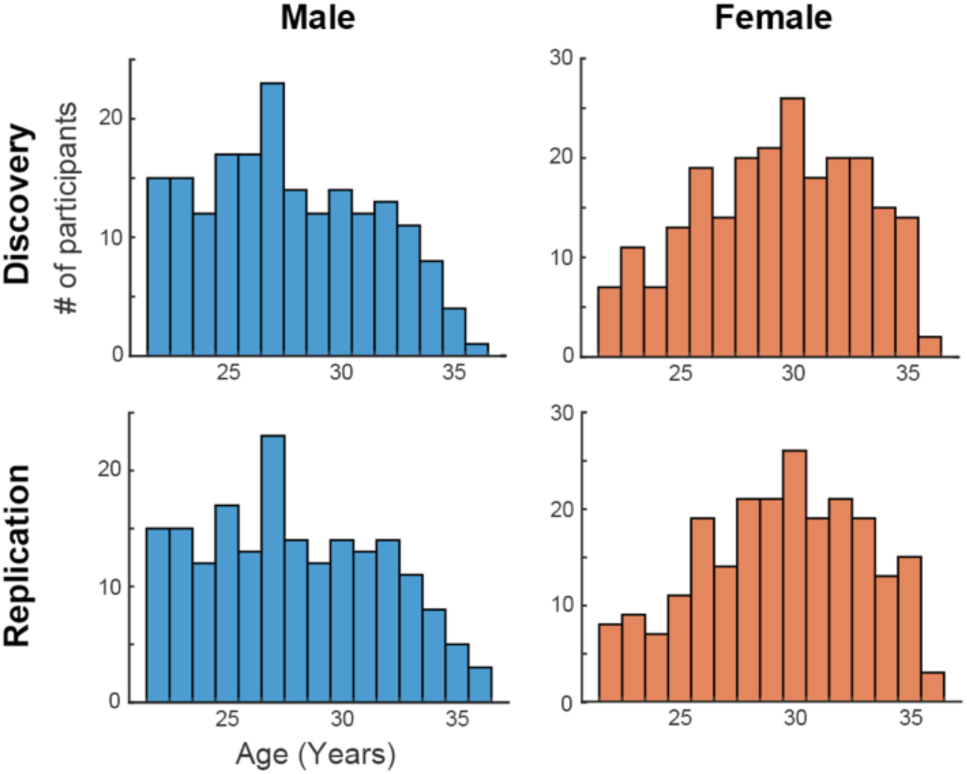
Histograms of age for male and female participants in discovery and replication groups.

### Experimental design

This study used two types of fMRI data from HCP: visual category localizer and resting-state data. Task design has been previously described in detail by Barch et al. (2013). The category localizer combines a visual category manipulation and a working memory (N-back) manipulation in a single task. Participants viewed images of either faces, whole bodies and body parts, hand-held tools, or scenes, organized into blocks by category. For brevity, the “bodies and body parts” condition will be referred to as “bodies.” Participants performed either a 0- or 2-back working memory task over individual images. Within each block, participants viewed a series of 10 images, each lasting 2s with a .5s inter-stimulus interval for a block duration of 25s. The start of each block also contained a 2.5s task cue. Each experimental run lasted 5:01 minutes (405 time points) and included 8 task blocks and 4 15s-long fixation blocks. Two runs of the task were acquired per participant. Resting-state fMRI data were acquired while participants had their eyes open and viewed a blank screen with a fixation cross. Four runs of resting-state data were collected, each lasting 1,200 time points.

### MRI data acquisition

MRI data were acquired on a custom Siemens 3T Skyra scanner, using previously described acquisition protocols (Van Essen et al., 2012). Brain anatomy was measured using high-resolution T1- and T2-weighted images. T1-weighted acquisitions used a 3D magnetization-prepared rapid acquisition gradient echo (*MPRAGE*) pulse sequence with inversion recovery, with parameters echo time (TE) = 2.14ms, repetition time (TR) = 2,400ms, inversion time (TI) = 1,000ms, field of view (FOV) = 180 x 224 x 224mm^3^, flip angle (FA) = 8°, bandwidth (BW) = 210Hz/pixel, echo spacing (ES) = 7.6ms, and voxel size = 0.7mm isotropic. T2-weighted acquisitions used a 3D SPACE sequence, with parameters TE = 565ms, TR = 3,200ms, FOV = 180 x 224 x 224mm^3^, variable FA, BW = 744Hz/pixel, ES 3.53ms, and voxel size = 0.7mm isotropic. Functional data acquisition used a gradient echo echo-planar-imaging sequence with parameters TE = 33.1ms, TR = 720ms, FOV = 208×180mm^2^, FA = 52°, BW = 2290Hz/pixel, ES = 0.58ms, voxel size = 2mm isotropic, and a multiband acceleration factor of 8.

### MRI data preprocessing

Data were preprocessed using the minimal preprocessing pipeline developed for the HCP, as described in detail elsewhere (Glasser et al., 2013). The pipeline uses tools from the FMRIB Software Library (FSL) and Freesurfer (Fischl, 2012; Smith et al., 2004). Both functional and anatomical images were corrected for gradient nonlinearity-induced distortion. Multiple anatomical acquisitions acquired within a given participant were registered and averaged. The average anatomical image was aligned to MNI152NLin6Asym space using a rigid-body transformation. Freesurfer’s recon-all was used to generate a mesh reconstruction of the cortical surface (Dale et al., 1999; Fischl, Sereno, & Dale, 1999). Individual surfaces were normalized to the fsaverage and fsLR template spaces using Freesurfer’s mris_register (Fischl, Sereno, Tootell, et al., 1999), followed by refinement using multimodal surface matching (Robinson et al., 2014).

For functional MRI data, motion parameters were first estimated using FSL’s MCFLIRT (Jenkinson et al., 2002). FSL’s topup was used to determine a nonlinear transformation to correct EPI distortion (Andersson et al., 2003). Rigid-body transformations to register functional to anatomical images were estimated using Freesurfer’s bbregister (Greve & Fischl, 2009). A single transformation combining correction for motion and distortion as well as anatomical registration was then applied to data with a single resampling. Independent component analysis (ICA-FIX) was used to remove noise signal, as implemented by the HCP pipeline (Salimi-Khorshidi et al., 2014). Lastly, data were resampled from the volume to cortical surface coordinates using ribbon-constrained mapping, and then transformed to 32k-density fsLR space. For region-of-interest definition, functional data were additionally spatially smoothed on the surface with a 4mm FWHM Gaussian kernel. All other analyses used unsmoothed data.

### General linear model (GLM)-based analysis

fMRI data analysis was performed using a custom pipeline (https://github.com/bmdeen/fmriPermPipe), with wrapper scripts used for this study available at (https://github.com/SocialMemoryLab/HCP_ATL). The analysis incorporates tools from multiple open-source software packages including FSL 6.0.7.4, AFNI, and Connectome Workbench 1.5. To estimate the strength of BOLD responses to each condition at each cortical surface coordinate, we first conducted a GLM-based analysis using AFNI’s 3dREMLfit. Regressors were defined as boxcar functions convolved with a canonical double gamma hemodynamic response function, for each of eight conditions: faces 0-back, bodies 0-back, tools 0-back, scenes 0-back, faces 2-back, bodies 2-back, tools 2-back, and scenes 2-back. Four contrasts were considered: faces vs other, bodies vs other, tools vs other, and scenes vs other. Statistics were corrected for temporal autocorrelation using an ARMA(1,1) model (Olszowy et al., 2019).

### Search space definition

To increase statistical sensitivity for studying regions with weak signal due to susceptibility artifacts, responses were analyzed using a region-of-interest (ROI)-based approach. Functional ROIs were defined within individual participants by applying functional criteria (described in ROI-based analysis: Discovery section) within anatomical search spaces. Four search spaces were considered: TP and PR, the focus of the study, as well as ventral occipitotemporal cortex (VOTC) and lateral occipitotemporal cortex (LOTC; Figure 2). VOTC and LOTC search spaces did not include parts of cortex expected to be scene-selective such as the parahippocampal gyrus, because we did not expect to observe scene responses in TP or PR.

**Figure 2:**
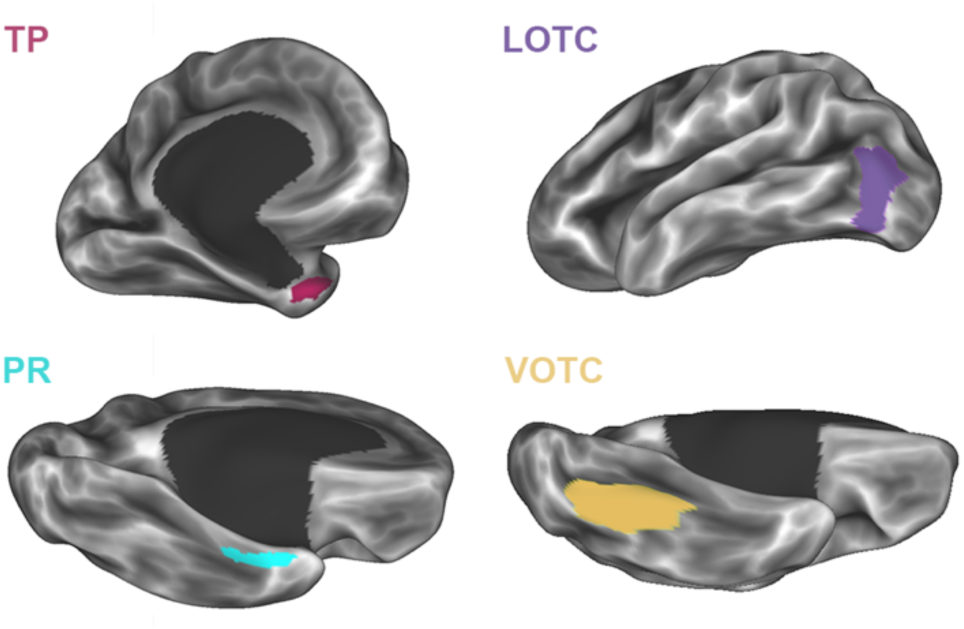
Location of anatomical search spaces used to define regions-of-interest. TP: temporal pole. PR: perirhinal cortex. LOTC: lateral occipitotemporal cortex. VOTC: ventral occipitotemporal cortex.

TP and PR were defined based on a probability map derived from hand-drawn regions in a separate set of participants (Deen et al., 2024). Borders were defined by anatomical landmarks previously established to correspond with cytoarchitectonic boundaries (Insausti et al., 1998). Binary mask images in fsLR space were averaged across participants to generate a probability map, smoothed with a 3mm FWHM Gaussian kernel, and thresholded at 60%. This threshold was selected to generate search spaces that were similar in size to hand-drawn TP and PR regions in individuals.

VOTC and LOTC were intended to cover parts of visual cortex that have been previously found to respond preferentially to specific categories, including faces, bodies, or inanimate objects. Functional ROIs defined with these search spaces were intended to provide a comparison to regions in TP and PR, and for the sake of assessing functional connectivity between anterior and posterior ROIs. Each search space was defined as the union of a set of regions from the HCP’s multimodal parcellation (Glasser et al., 2016). VOTC included areas VVC, V8, and FFC. LOTC included PIT, LO1, LO2, LO3, V4t, and MT.

### ROI-based analysis: Discovery

ROI-based analysis was conducted using a two-step procedure. We first searched for dissociable patterns of response across categories in the discovery group, and then confirmed these patterns in the replication group. To avoid imposing any prior hypotheses about which categories would drive responses in TP and PR, we considered all possible comparisons of one image category to the others: faces vs other, bodies vs other, tools vs other, and scenes vs other. ROIs were defined separately for left and right TP and PR, leading to 4*4 = 16 functional ROIs (four search spaces by four contrasts). ROIs were defined as the set of surface coordinates with the top *M*% of statistical values for a given contrast within a given search space. Relative to using a fixed statistical threshold, this approach has the benefit of defining ROIs in all participants for all contrasts, and ensuring consistency of ROI size across participants. For main analyses the value *M* = 5 was used, based on our prior work (Deen et al., 2024), while additional analyses used a range of values from *M* = 5 to 50 to determine how results were influenced by ROI size. To avoid circularity in ROI definition and response extraction, independent data were used for each: we first defined ROIs in run 1 and extracted responses in run 2; then defined ROIs in run 2 and extracted responses in run 1; and finally averaged extracted responses across runs. Spatially smoothed data were used to define ROIs, while unsmoothed data were used to extract responses. Given our interest in visual category preferences, responses were averaged across 0-and 2-back conditions for statistical analysis and visualization.

From the resulting set of responses from 16 ROIs, we identified a set of distinct response profiles to study further. For a given search space, response profiles from two contrasts were considered distinct if they differed in the *order* of response strengths across the four categories (e.g. faces > bodies > tools > scenes). This resulted in a set of three distinct response profiles in each hemisphere – one in TP, and two in PR. Among ROIs that were deemed not distinct by this criterion, response magnitudes across conditions did not differ substantially. Given these results, we selected three contrasts with distinct responses to use for further investigation: faces vs other in TP and PR, and tools versus other in PR. The ROIs defined by these contrasts will be denoted face-preferring and tool-preferring, irrespective of their observed response profile, to specify how the ROI was defined.

### ROI-based analysis: Replication

We next ran ROI analyses in the replication group. ROI definition and response extraction were performed as described above for the discovery group. Responses to each pair of conditions for each ROI were compared using a linear mixed model implemented by MATLAB’s fitlme, with each run treated as a separate observation, including a random intercept term for participant. All comparisons were thresholded at P < .017 (0.05/3) two-tailed, applying Bonferroni correction across the three ROI-defining contrasts. To determine whether response profiles differed significantly across ROIs, we ran a mixed effects ANOVA to test for an ROI by condition by hemisphere interaction, including a random intercept term for participant. Additionally, to determine whether effects of hemisphere would emerge with a more sensitive model, we ran mixed effects ANOVAs for each ROI testing for a condition by hemisphere interaction, including a random intercept term for participant. To qualitatively compare responses in TP and PR to well-characterized regions in VOTC and LOTC, we conducted ROI analyses in the latter search spaces, defining ROIs using faces > other and tools > other contrasts. Because category-sensitive responses in VOTC and LOTC have been extensively documented in prior research, statistics were not performed on ROI responses.

### Spatial position analysis

Prior research has documented systematic spatial relationships between occipitotemporal regions with differing category preferences. We next asked whether any systematic spatial relationship exists for face- and tool-preferring regions with PR. In each participant, we computed the mean volumetric coordinates of ROIs in PR (along with VOTC and LOTC for comparison), and then computed the difference in spatial position between face- and tool-preferring ROIs along the x, y, and z axes. Paired two-sample, two-tailed *t*-tests were used to statistically assess differences in spatial position between the two types of ROI.

### Functional connectivity analysis

We next asked whether ROIs in ATL could be differentiated based on their profile of resting-state functional connectivity to other cortical regions. All analyses were performed separately in the discovery and replication groups, to evaluate consistency and provide an internal replication. Functional connectivity analyses were conducted with three complementary approaches: 1) whole-brain, 2) functional network-based, and 3) ROI-based. For whole brain analyses, time series correlations were computed between the spatially averaged signal from each functional ROI and signal from each other surface coordinate. Correlation maps were then averaged across participants.

To assess functional connectivity of ATL subregions with specific functional networks, we defined network parcellations of cortex in individual participants using the multi-session hierarchical Bayesian model (MSHBM) algorithm (Kong et al., 2019). As a group-level prior, we used a 15-network parcellation that has previously been demonstrated to identify meaningful and functionally distinct systems (Du et al., 2024). Networks included default A (DN-A), default B (DN-B), language (LANG), frontoparietal A (FPN-A), frontoparietal B (FPN-B), salience / parietal memory network (SAL/PMN), cingulo-opercular (CG-OP), dorsal attention A (dATN-A), dorsal attention B (dATN-B), premotor-posterior parietal (PM-PPr), somatomotor A (SMOT-A), somatomotor B (SMOT-B), auditory (AUD), visual-central (VIS-C), and visual-peripheral (VIS-P). Time series correlations were computed between spatially averaged signal from ATL ROIs and each network. To test for statistical differences in ROIs’ patterns of functional connectivity, we used a linear mixed effects model with an ROI x network interaction term, including a random intercept for participant.

Lastly, we asked whether face- and tool-preferring ATL ROIs had stronger functional connectivity to regions of occipitotemporal cortex with like category preferences. We computed time series correlations between spatially averaged signal from ATL ROIs with LOTC and VOTC ROIs. We focused on within-hemisphere correlations (e.g. left PR to left LOTC and VOTC). Differences between correlations for face-preferring and tool-preferring OTC regions were assessed using paired two-sample, one-tailed *t*-tests.

## Results

### ATL contains three distinct profiles of visual category preference

Using category localizer data from the Human Connectome Project, we first searched for distinct patterns of category preference within the ATL in a subset of participants termed the discovery group. Within anatomical search spaces covering TP and PR, we defined functional ROIs as the set of surface coordinates with the strongest preference for each category (faces, bodies, tools, or scenes) relative to the others, and then measured each ROI’s response profile in independent data (Figure 3).

**Figure 3:**
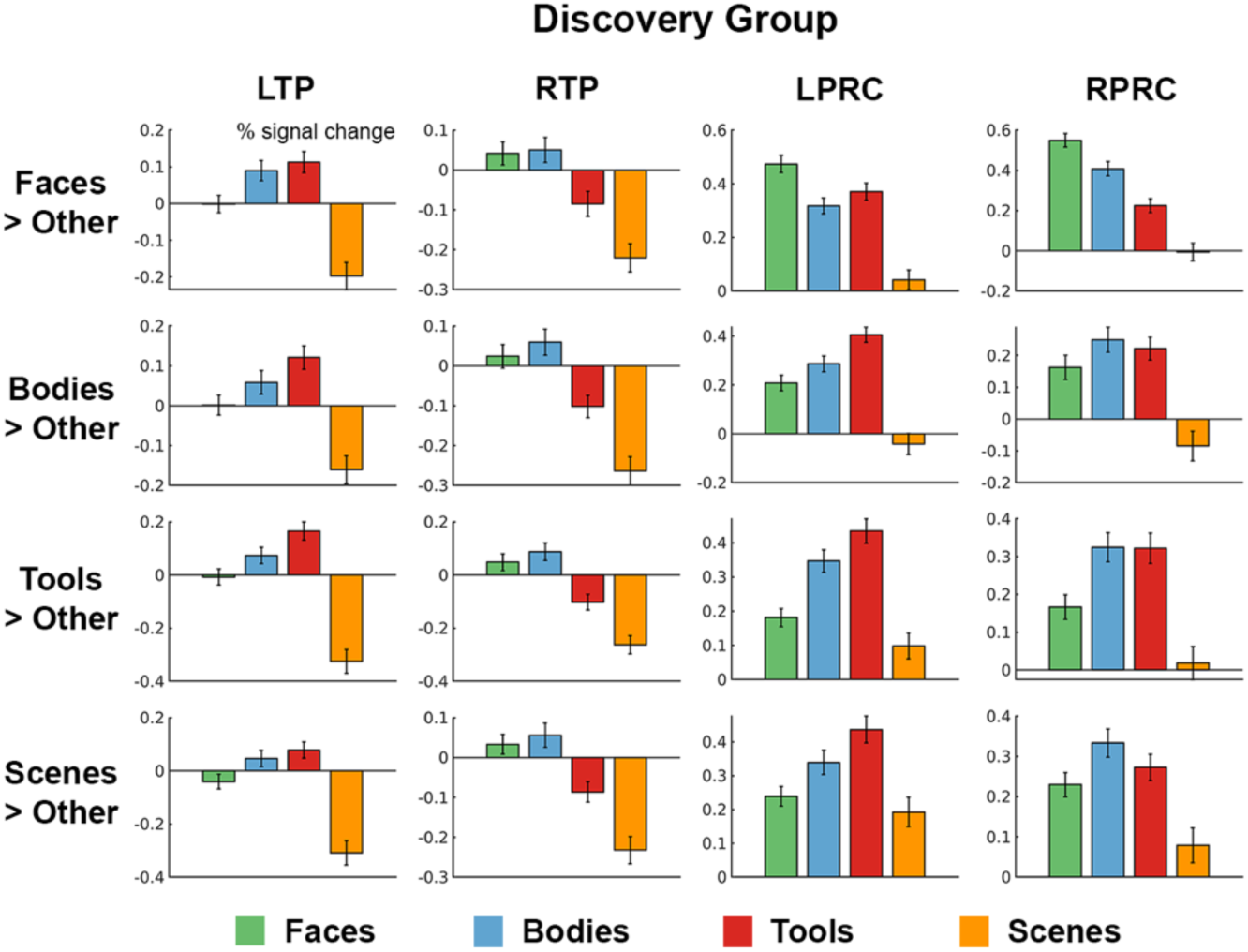
Region-of-interest (ROI)-based analysis reveals three patterns of category preference within the anterior temporal lobe. Responses (% signal change) in the discovery group from functional ROIs defined using four contrasts (each category compared to the others) in four anatomical search spaces. Error bars reflect standard error across participants.

In TP, similar response profiles were obtained regardless of the contrast used define the ROI. This profile was characterized by responses near baseline to the object conditions (faces, bodies, and tools), and a decreased response to the scene condition. The left hemisphere showed a stronger preference for bodies and tools, while the right hemisphere showed a stronger preference for faces and bodies. This result indicates visual responses within TP in a given hemisphere in this dataset are largely homogenous across cortical location.

By contrast, PR showed a strikingly different pattern, with object conditions eliciting responses above baseline. Across the four ROI-defining contrasts, two distinct profiles were observed in PR: the faces vs other contrast yielded a preference for faces over other conditions, while the other three contrasts (bodies vs other, tools vs other, and scenes vs other) yielded a preference for bodies and tools over other conditions. Taken together, these indicate that the ATL contains at least three distinct patterns of visual response across categories.

### Three response profiles replicate in independent data

To replicate and statistically evaluate these results, we next conducted ROI analyses in a separate subset of participants termed the replication group. Based on the discovery group results, we focused on regions defined by three contrasts: faces vs other in TP, faces vs other in PR, and tools vs other in PR. The resulting ROIs will be termed face-preferring TP (fTP), face-preferring PR (fPR), and tool-preferring PR (tPR), although we emphasize that this language is used only to specify the contrast used to define ROIs, and not the resulting response profile or an interpretation of each area’s function.

Response profile in the replication group were highly similar to the discovery group, with three distinct patterns observed in each hemisphere (Figure 4). To determine whether responses differed significantly across ROIs, we conducted a mixed effects ANOVA testing for an ROI x condition x hemisphere interaction. This identified significant effects of ROI (*F*(2,9936) = 47.56, *P* < 10^-20^) and condition (*F*(3,9936) = 20.08, *P* < 10^-12^), and an ROI by condition interaction (*F*(6,9936) = 8.15, *P* < 10^-8^). The model did not show significant effects of hemisphere (*F*(1,9936) = .17, *P* = .68) or interaction terms for ROI by hemisphere (*F*(2,9936) = .60, *P* = .55), condition by hemisphere (*F*(3,9936) = 2.44, *P* = .06), or ROI by condition by hemisphere (*F*(6,9936) = .27, *P* = .95). These results demonstrate that patterns of visual response across conditions differ significantly across the three functional ROIs. To determine whether effects of hemisphere would be observed with a more sensitive test, we next conducted mixed effects ANOVAs for each ROI separately, testing for a condition by hemisphere interaction. Effects of condition were observed for all three ROIs (fTP: *F*(3,3312) = 28.32, *P* < 10^-17^; fPR: *F*(3,3312) = 38.42, *P* < 10^-23^; tPR: *F*(3,3312) = 16.65, *P* < 10^-10^). Effects of hemisphere were not observed for any ROI (fTP: *F*(1,3312) = .24, *P* = .62; fPR: *F*(1,3312) = 1.44, *P* = .23; tPR: *F*(1,3312) = .03, *P* = .86). An interaction between condition and hemisphere was observed for fTP (*F*(3,3312) = 3.45, *P* < .0166 = .05/3), but not fPR (*F*(3,3312) = .91, *P* = .44) or tPR (*F*(3,3312) = 1.51, *P* = .21). These results demonstrate that the response of TP across visual categories differs significantly by hemisphere.

**Figure 4.**
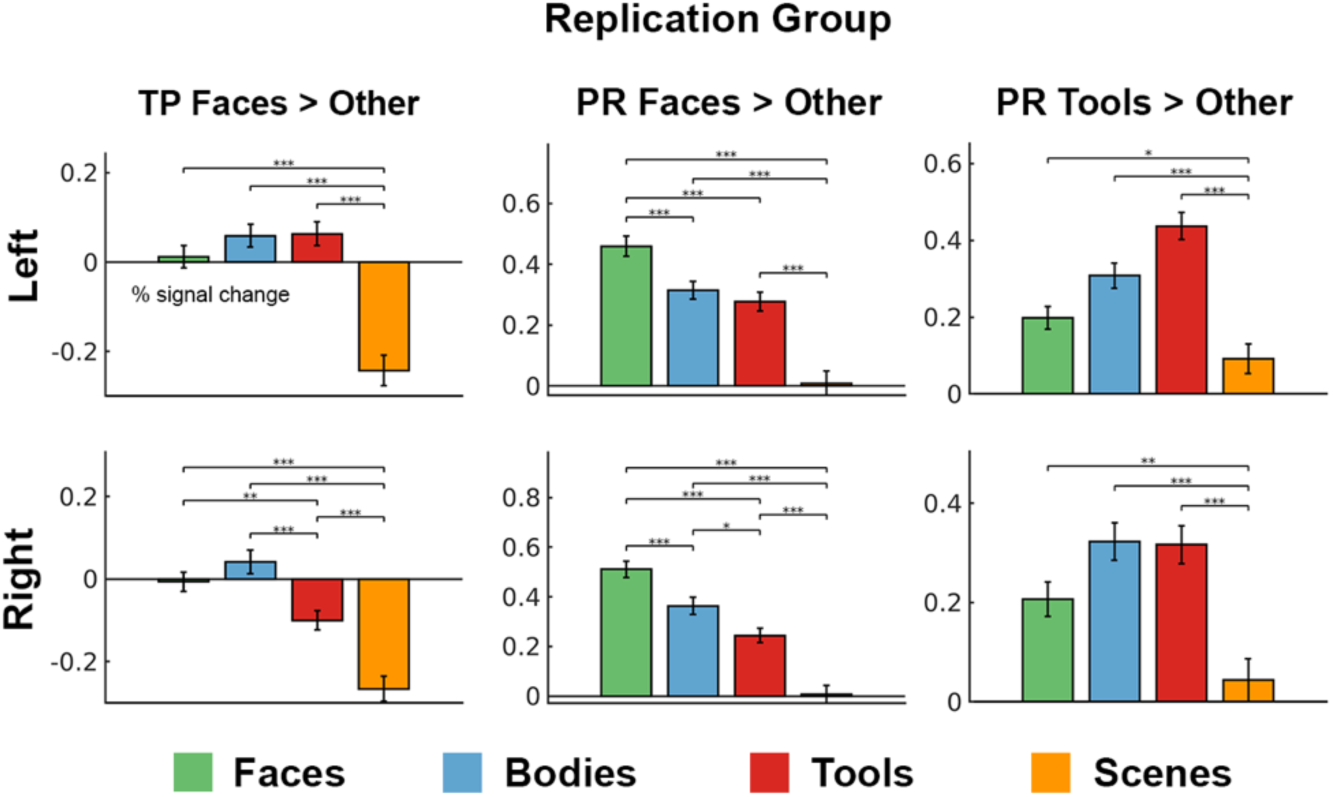
Three distinct response profiles replicate in an independent sample of participants. Responses (% signal change) in the replication group from ROIs defined by the contrast faces > other (TP and PR) and tools > other (PR). Error bars reflect standard error across participants. * *P* < .017 = .05/3, ** *P* < .001, *** *P* < 10^-4^.

We next tested for differences in response between each pair of conditions within each ROI. Left fTP responded preferentially to faces (*t*(829) = 8.64, P < 10^-16^), bodies (*t*(829) = 7.26, P < 10^-12^), and tools (*t*(829) = 6.73, P < 10^-10^) over scenes. Right fTP responded preferentially to faces (*t*(829) = 7.93, P < 10^-14^), bodies (*t*(829) = 6.70, P < 10^-10^), and tools (*t*(829) = 4.97, P < 10^-6^) over scenes, as well as faces (*t*(829) = 3.45, P < .001) and bodies (*t*(829) = 4.04, P < 10^-4^) over tools. These results suggest that left TP has a preference for objects over scenes, while right TP has a preference for social objects (faces and bodies) over nonsocial objects and scenes.

Left fPR responded preferentially to faces over bodies (*t*(829) = 4.62, P < 10^-5^), tools (*t*(829) = 4.75, P < 10^-5^), and scenes (*t*(829) = 12.92, P < 10^-34^), as well as to bodies (*t*(829) = 6.75, P < 10^-10^) and tools (*t*(829) = 5.55, P < 10^-7^) over scenes. Right fPR responded preferentially to faces over bodies (*t*(829) = 4.28, P < 10^-4^), tools (*t*(829) = 7.22, P < 10^-11^), and scenes (*t*(829) = 12.93, P < 10^-34^), as well as to bodies (*t*(829) = 6.85, P < 10^-10^) and tools (*t*(829) = 6.30, P < 10^-9^) over scenes, and bodies over tools (*t*(829) = 3.08, P < .005). These results suggest that PR contains a face-sensitive functional subregion with a preference for faces over bodies over tools over scenes.

Left tPR responded preferentially to tools (*t*(829) = 6.35, P < 10^-9^), bodies (*t*(829) = 4.48, P < 10^-5^), and faces (*t*(829) = 2.76, P < .01) over scenes. Right tPR responded preferentially to tools (*t*(829) = 5.52, P < 10^-7^), bodies (*t*(829) = 4.72, P < 10^-5^), and faces (*t*(829) = 3.74, P < .001) over scenes. These results suggest that in addition to a face-preferring subregion, PR contains a distinct functional subregion that responds generally to objects over scenes.

Do responses in TP and PR depend on the size criterion used to define ROIs? While the above analyses define ROIs as surface coordinates with the top 5% of statistics for a given contrast, we next varied this value from 5 – 50%. We then computed the value of the ROI-defining contrasts (faces vs other or tools vs other) in independent data, and plotted this as a function of ROI size (Figure 5). Contrast values in fTP did not vary substantially by ROI size, consistent with the observation that task responses do not vary by cortical location within TP. For fPR and tPR, contrast values decreased with increasing ROI size, but remained greater than zero across the range of sizes tested. This indicates that our results do not depend strongly on the size criterion used to define ROIs.

**Figure 5.**
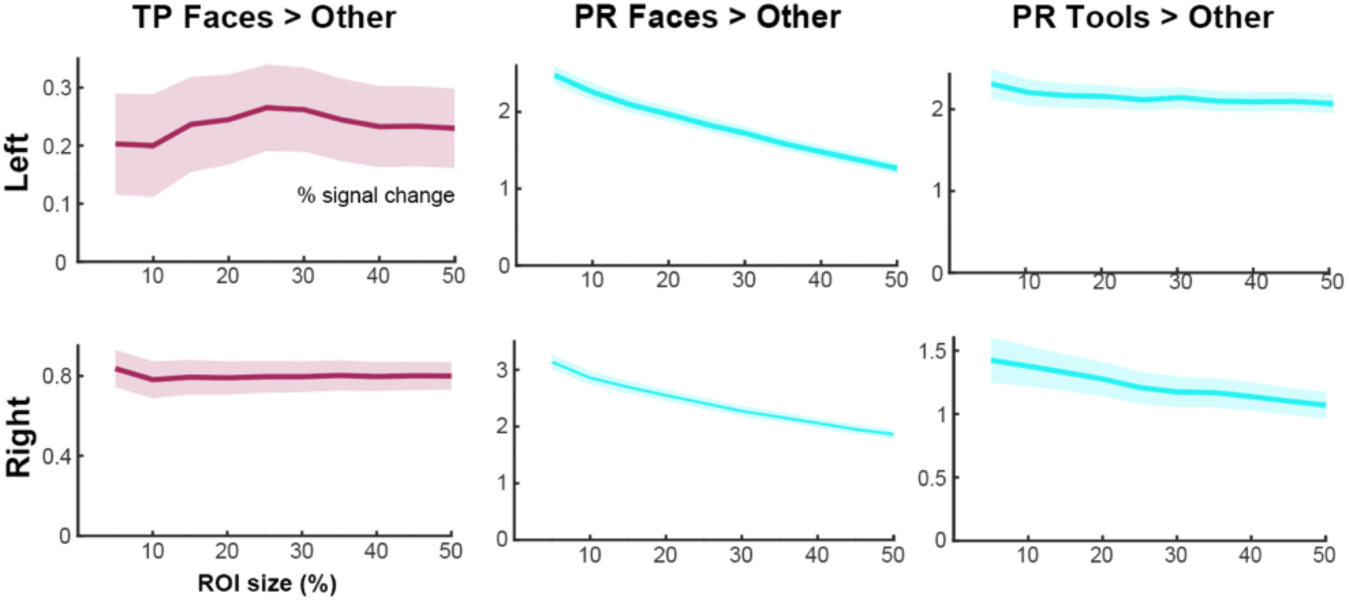
Category preferences in the anterior temporal lobe are consistent across ROI size. Contrast value (faces > other or tools > other expressed in % signal change) plotted as a function of ROI size (% of coordinates within a search space). Shaded regions reflect standard error across participants.

How do visual responses in TP and PR compare to previously established category-sensitive regions in posterior occipitotemporal cortex? To ask this question, we measured responses in face- and tool-preferring ROIs in VOTC and LOTC (Figure 6). Face-preferring ROIs across both VOTC and LOTC showed the strongest response to faces, followed by bodies, tools, and scenes, similar to the pattern observed for face-preferring PR. Tool-preferring ROIs in LOTC showed a strong response to tool and body conditions, and a weaker response to faces and scenes. Tool-preferring ROIs in VOTC showed the strongest response to tools, then bodies and scenes, then faces. These response profiles are similar to what we observed for tool-preferring PR, which showed the strongest responses to tools and bodies. These results show that commonly reported category-sensitive responses in occipitotemporal cortex can be reproduced with this dataset and analysis approach. Furthermore, they indicate that patterns of category preference observed in posterior occipitotemporal cortex are largely reproduced in perirhinal cortex.

**Figure 6.**
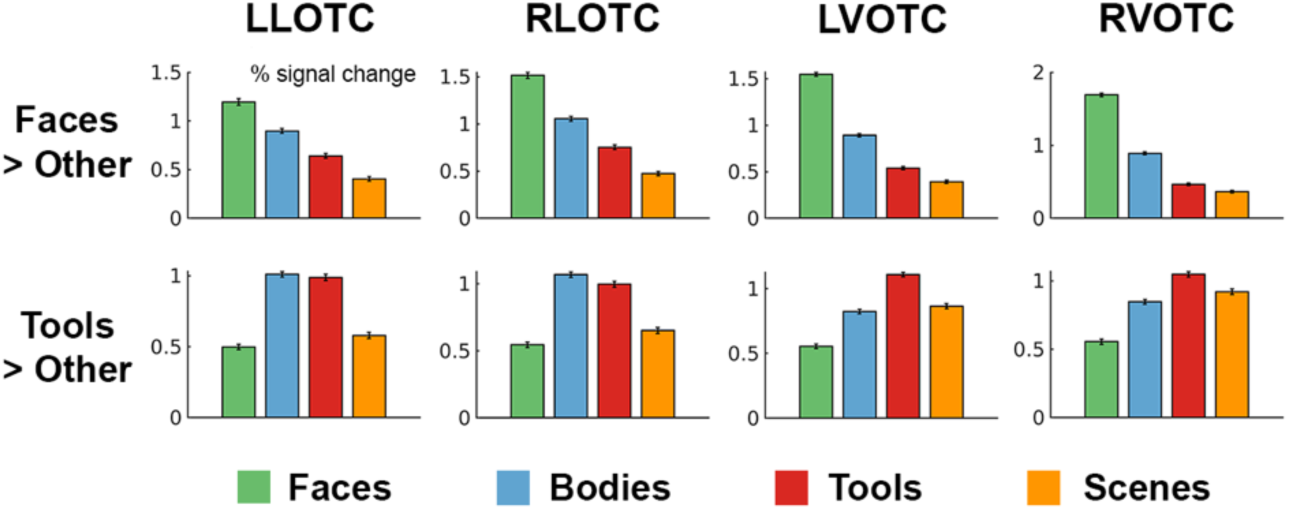
ROIs in lateral occipitotemporal cortex (LOTC) and ventral occipitotemporal cortex (VOTC) show established patterns of face and object preference. Responses (% signal change) from functional ROIs defined by faces > other and tools > other contrasts within LOTC and VOTC search spaces. Error bars reflect standard error across participants.

### Face and tool responses in PR lack a reliably distinct spatial position

Do face- and tool-preferring ROIs in PR differ in spatial location? Within LOTC and VOTC, prior work has shown that these responses have systematic spatial relationships. In LOTC, face responses in the occipital face area are located inferior to tool responses in the posterior middle temporal gyrus (Bracci et al., 2012; Chao et al., 1999; Pillet et al., 2024). In VOTC, face responses are located lateral to tool responses on the fusiform gyrus (Chao et al., 1999; Mahon et al., 2007; Noppeney et al., 2006). To evaluate spatial position, we computed mean x-, y-, and z-coordinates of face- and tool-preferring ROIs from the replication group in data rigidly aligned with MNI space, computed the difference between positions of face- and tool-preferring ROIs in each individual participant, and assessed the distribution of these difference values across participants.

Violin plots of spatial difference values are shown in Figure 7. We first describe results for LOTC and VOTC, to validate our approach based on predictions from prior work. For left LOTC, face responses were significantly lateral (*t*(414) = –2.61, *P* < .01, Cohen’s *d* = –.13, mean spatial distance Δ = –.7mm), anterior (*t*(414) = 5.61, *P* < 10^-7^, *d* = .28, Δ = 1.6mm), and inferior (*t*(414) = –10.80, *P* < 10^-23^, *d* = –.53, Δ = –5.4mm) to tool responses. For right LOTC, face responses were significantly lateral (*t*(414) = 3.85, *P* < .001, *d* = .19, Δ = 1.1mm) and inferior (*t*(414) = – 16.30, *P* < 10^-45^, *d* = –.80, Δ = –7.5mm) to tool responses, with no difference observed in the anterior-posterior direction (*P* > .3). These results replicate prior work finding face responses inferior to tool responses in LOTC and suggest that they are also positioned slightly laterally.

**Figure 7.**
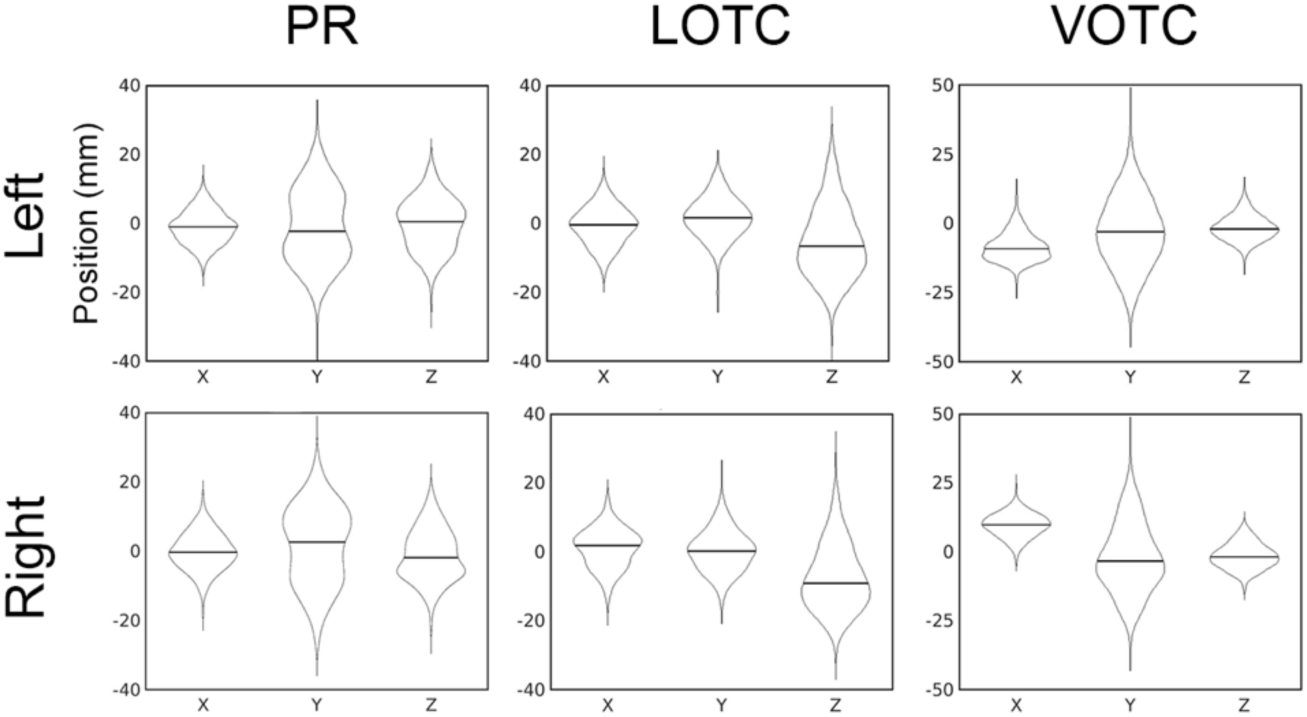
Face- and tool-preferring ROIs in PR do not show systematic differences in spatial position, unlike LOTC and VOTC. Violin plots across participants of the differences in position between face- and tool-preferring ROIs. Coordinates are aligned with MNI152 space and in RAS orientation.

For left VOTC, face responses were significantly lateral (*t*(414) = –33.92, *P* < 10^-120^, *d* = –1.67, Δ = –8.5mm), posterior (*t*(414) = –4.74, *P* < 10^-5^, *d* = –.23, Δ = –2.9mm), and inferior (*t*(414) = –7.13, *P* < 10^-11^, *d* = –.35, Δ = –1.6mm) to tool responses. For right LOTC, face responses were significantly lateral (*t*(414) = 46.37, *P* < 10^-165^, *d* = 2.28, Δ = 9.7mm), posterior (*t*(414) = –3.65, *P* < .001, *d* = –.18, Δ = –2.2mm), and inferior (*t*(414) = –6.91, *P* < 10^-10^, *d* = –.34, Δ = –1.5mm) to tool responses. These results replicate prior work finding face responses are positioned medially to tool responses within VOTC, and suggest that they are also positioned slightly posteriorly and inferiorly.

For left PR, face responses were significantly lateral (*t*(414) = –4.80, *P* < 10^-5^, *d* = –.24, Δ = 1.1mm) to tool responses, showing no difference in the anterior-posterior (*P* > .05) or superior-inferior (*P* > .6) dimensions. For right PR, face responses were significantly anterior (*t*(414) = 3.09, *P* < .01, *d* = –.15, Δ = 1.6mm) and inferior (*t*(414) = –3.43 *P* < .001, *d* = –.17, Δ = 1.2mm) to tool responses, with no difference in the medial-lateral dimension (*P* > .7). Given the small effect sizes, small spatial distances, and inconsistent results across hemispheres, we conclude that there are no clear differences in spatial position between face- and tool-preferring ROIs in PR, unlike LOTC and VOTC. This indicates that category preferences in the ATL may have a patchy or “salt-and-pepper” spatial organization, distinct from the well-separated regions observed in posterior occipitotemporal cortex.

### ATL subregions have dissociable functional connectivity patterns

Do ATL subregions with distinct visual responses also show differing patterns of functional connectivity to the rest of cortex? We ask this question by measuring time series correlations in resting-state data. We use multiple approaches with complementary strengths: 1) whole-cortex, measuring correlations from functional ROIs all other cortical surface coordinates; 2) network-based, measuring correlations to specific functional networks derived from an individualized cortical parcellation; and 3) ROI-based, measuring correlations to face- and tool-preferring ROIs in LOTC and VOTC.

Whole-cortex patterns of functional connectivity from each ATL ROI are shown in Figures 8 (discovery group) and 9 (replication group). For ease of visualization given our focus on differences between ROIs, only within-hemisphere correlations are shown; between hemisphere correlations were generally weaker but similar in spatial pattern. Results were highly consistent across the discovery and replication groups and left and right hemispheres and will therefore be described collectively. A clear difference was observed between functional connectivity between face-preferring TP and the PR ROIs. fTP showed strongest correlations with areas associated with the default mode network, including medial parietal cortex, medial prefrontal cortex, temporo-parietal junction, anterior superior temporal sulcus, inferior frontal gyrus, and superior frontal sulcus. By contrast, fPR and tPR showed strongest correlations with areas associated with visual processing, including ventral occipitotemporal cortex, lateral occipitotemporal cortex, the intraparietal sulcus, and regions of lateral prefrontal cortex resembling components of the dorsal attention network. While functional connectivity maps for fPR and tPR were similar in large-scale patterns, several differences were observed reliably across groups and hemispheres. fPR showed stronger correlations to areas associated with face processing, including posterior and anterior superior temporal sulcus, and the lateral fusiform gyrus.

**Figure 8.**
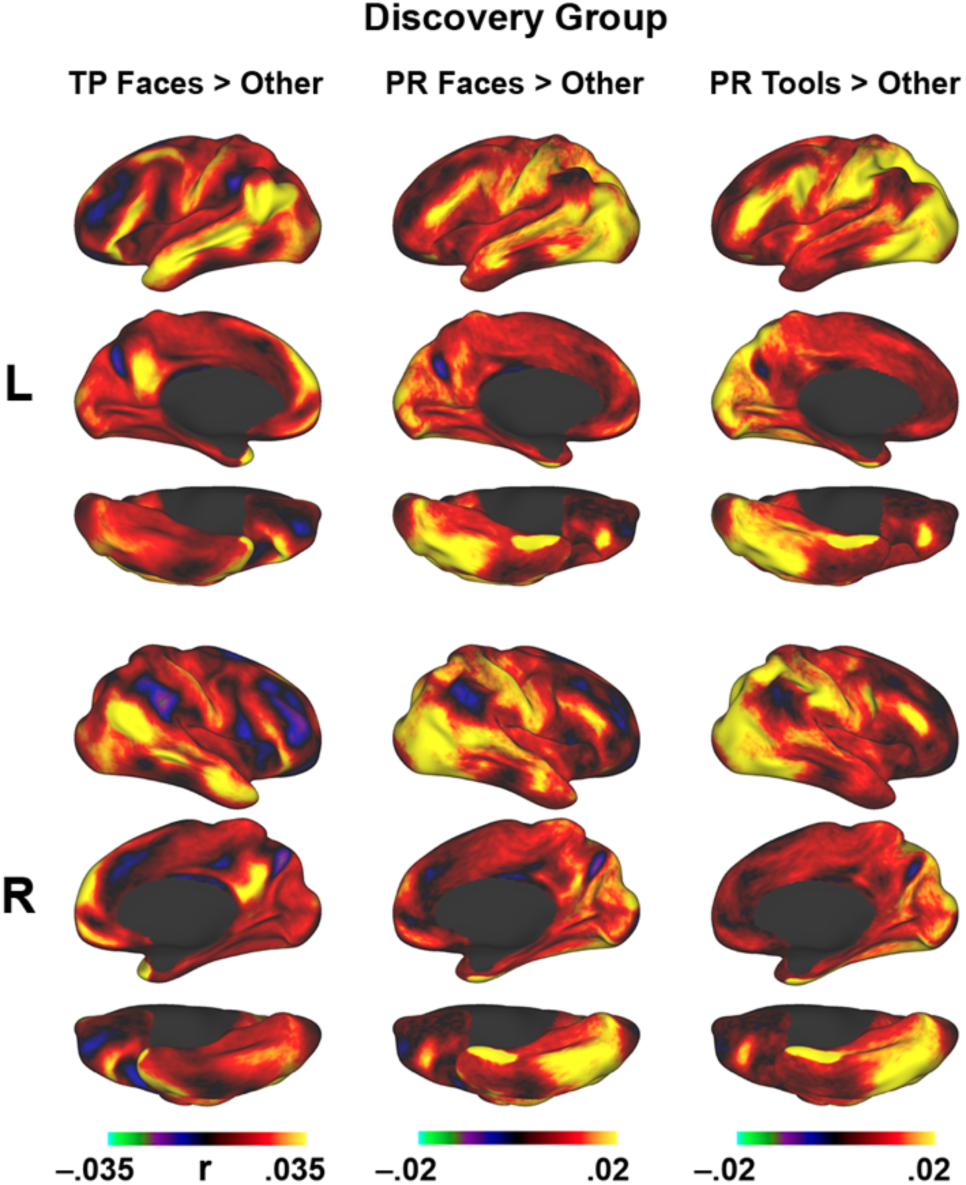
Anterior temporal lobe subregions have dissociable patterns of cortex-wide functional connectivity. Maps of resting-state correlations from functional ROI seed regions in the discovery group. L = left hemisphere; R = right hemisphere.

**Figure 9.**
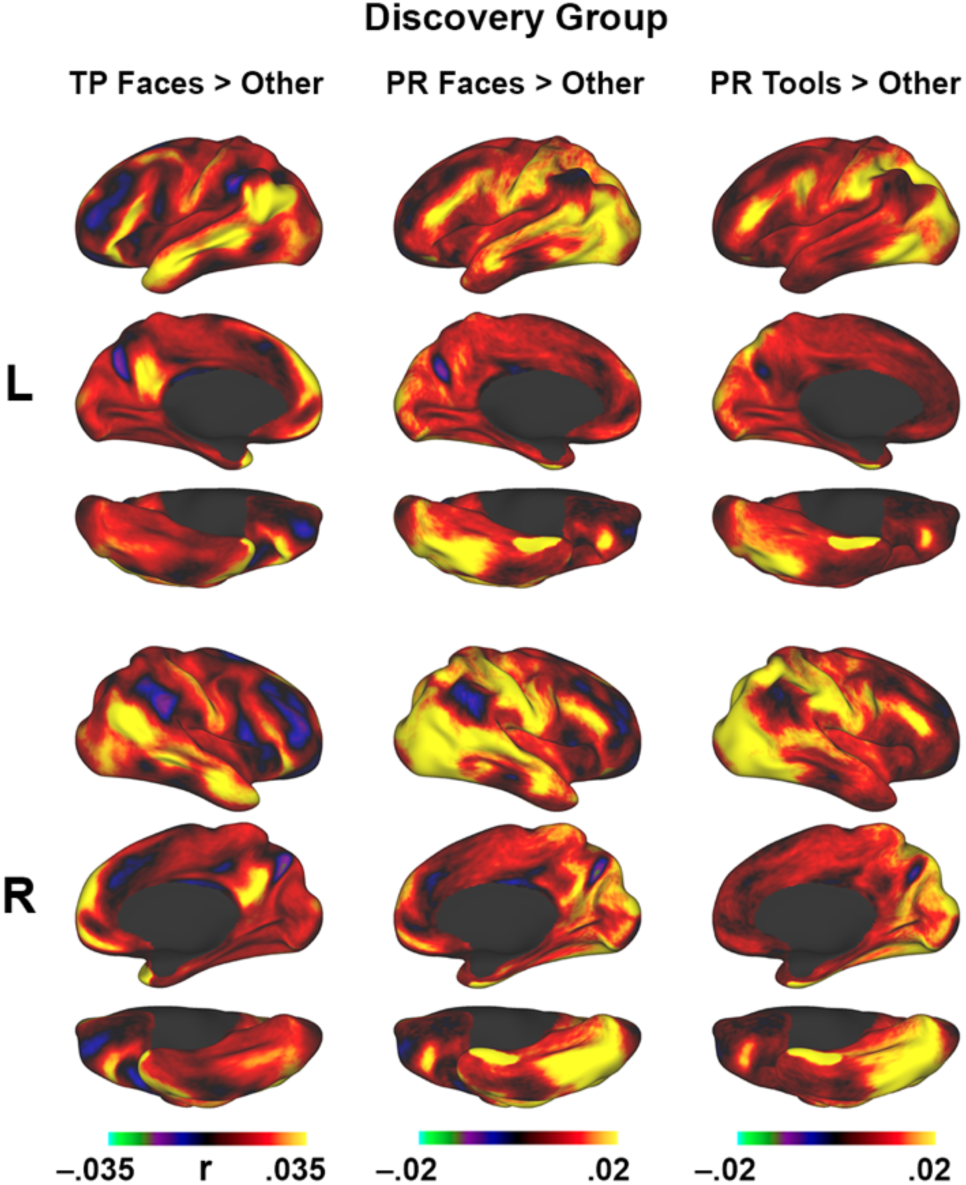
Dissociable patterns of functional connectivity across anterior temporal lobe subregions replicate in an independent sample. Maps of resting-state correlations from functional ROI seed regions in the discovery group. L = left hemisphere; R = right hemisphere.

To determine how these differences relate to functional connectivity with specific functional networks, and to assess the differences statistically, we next evaluated functional connectivity between ATL ROIs and fifteen individually defined networks (Figure 10). Results were highly consistent across the discovery and replication groups and left and right hemispheres and will therefore be described collectively. Overall, correlation strengths were relatively low, with maximum correlations (cross-subject mean) around .1 for TP and .07 for PR. These low values are not surprising given 1) the low signal strength expected in regions of the ATL; and 2) the emphasis of the HCP data acquisition approach on resolution over signal strength, using small voxels and a high multiband acceleration factor. Nevertheless, correlations showed systematic differences in their pattern across networks.

**Figure 10.**
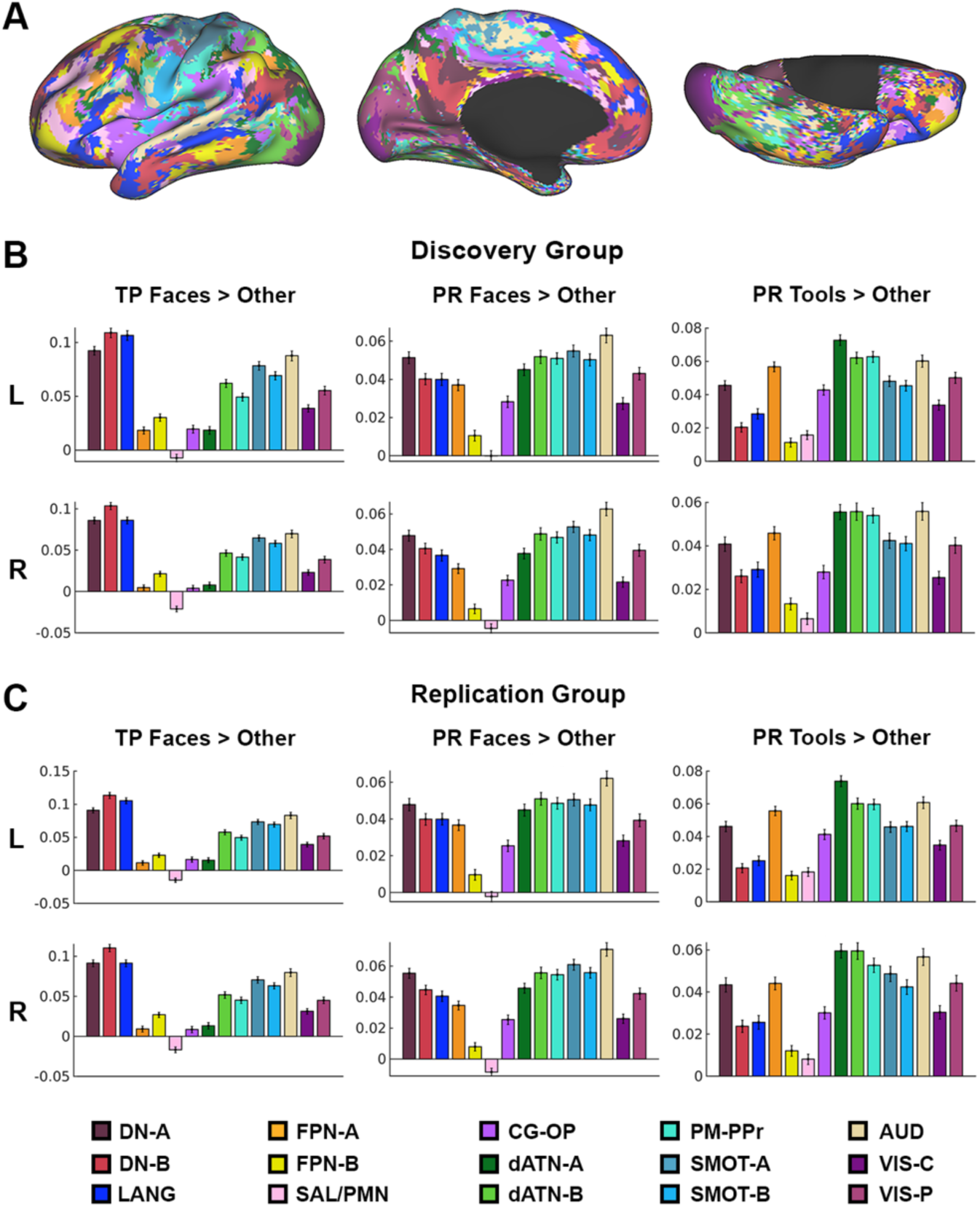
Anterior temporal lobe subregions have dissociable patterns of functional connectivity to individualized functional networks. (A) Parcellation of cortex into 15 functional networks in an example participant. (B-C) Resting-state correlations between functional ROIs and each network in the discovery (B) and replication groups (C). L = left hemisphere; R = right hemisphere. Error bars reflect standard error across participants.

Consistent with the whole-cortex results, fTP was most strongly correlated with DN-B, followed by DN-A and LANG. fPR and tPR showed a more diffuse pattern of correlations across networks, including strongest correlations with dATN-A, dATN-B, PM-PPr, SMOT-A, SMOT-B, and AUD. fPR was most strongly correlated with AUD. While this may appear surprising given this network’s presumed auditory function, the network in individual participants contains much of the upper back of the superior temporal sulcus, where correlations with fPR were observed in the whole-cortex analysis. tPR was most strongly correlated with dATN-A, dATN-B, and PM-PPr.

Do patterns of functional connectivity to different networks vary significantly by ROI? We tested this using a linear mixed effects model assessing the interaction between ROI and network. In the discovery group, we observed significant main effects of ROI (*F*(2,37305) = 129.29, *P* < 10^-55^) and network (*F*(14,37305) = 263.23, *P* < 10^-300^) and an interaction between ROI and network (*F*(28,37305) = 78.85, *P* < 10^-300^). In the replication group, we observed significant main effects of ROI (*F*(2,37305) = 129.62, *P* < 10^-56^) and network (*F*(14,37305) = 280.61, *P* < 10^-300^) and an interaction between ROI and network (*F*(28,37305) = 85.61, *P* < 10^-300^). These results demonstrate that fTP, fPR, and tPR ROIs have significantly different patterns of functional connectivity across networks.

Lastly, we ask whether ATL regions are preferentially functionally connected with occipitotemporal regions that have like visual category preferences (Figure 11). In the discovery group, left fTP was more correlated with face- vs tool-preferring VOTC (*t*(414) = 6.06, *P* < 10^-8^), while a significant difference was not observed for LOTC (*t*(414) = 1.23, *P* = .11). Right fTP was more correlated with face- vs tool-preferring VOTC (*t*(414) = 6.95, *P* < 10^-16^) while the effect for LOTC did not reach significance after multiple comparison correction (*t*(414) = 2.09, *P* = .019). Left fPR was more correlated with face- vs tool-preferring LOTC (*t*(414) = 2.75, *P* < .005) and with face- vs tool-preferring VOTC (*t*(414) = 5.81, *P* < 10^-8^). Right fPR was more correlated with face- vs tool-preferring LOTC (*t*(414) = 2.85, *P* < .005) and with face- vs tool-preferring VOTC (*t*(414) = 6.85, *P* < 10^-10^). Left tPR was more correlated with tool- vs face-preferring LOTC (*t*(414) = 4.29, *P* < 10^-5^) while a significant difference was not observed for VOTC (*t*(414) = 1.62, *P* = .053). Right tPR was more correlated with tool- vs face-preferring LOTC (*t*(414) = 2.44, *P* < .01), while the effect for VOTC did not reach significance after multiple comparison correction (*t*(414) = 2.09, *P* = .019).

**Figure 11.**
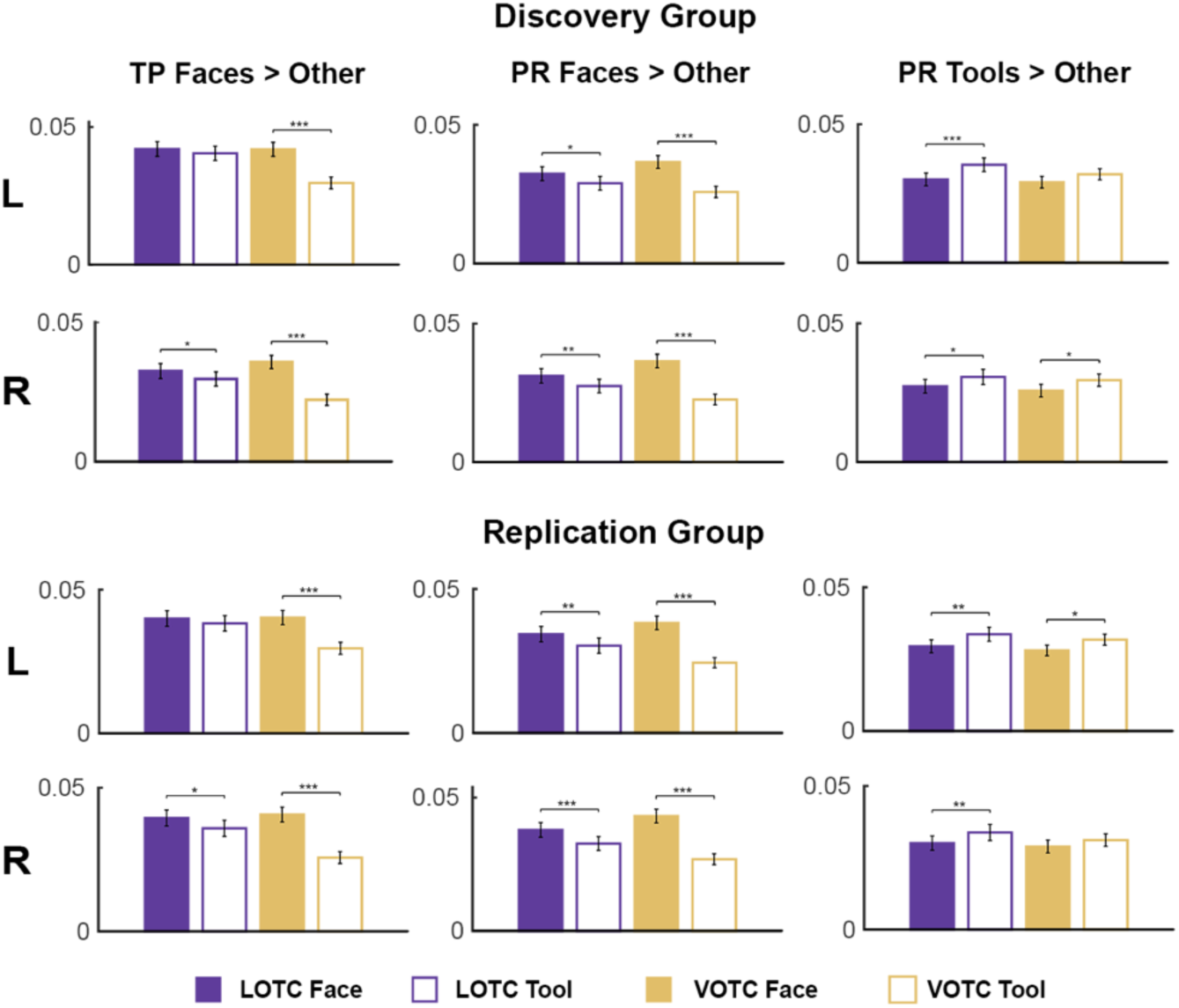
Face- and tool-preferring subregions of anterior temporal cortex are functionally connected with occipotemporal regions with like category preferences. Resting-state correlations between ROIs in TP/PR and LOTC/VOTC. L = left hemisphere, R = right hemisphere. Error bars reflect standard error across participants. * *P* < .017 = .05/3, ** *P* < .001, *** *P* < 10^-4^.

In the replication group, left fTP was more correlated with face- vs tool-preferring VOTC (*t*(414) = 6.47, *P* < 10^-9^), while a significant difference was not observed for LOTC (*t*(414) = 1.13, *P* = .13). Right fTP was more correlated with face- vs tool-preferring LOTC (*t*(414) = 2.25, *P* < .017 = .05/3) and with face- vs tool-preferring VOTC (*t*(414) = 8.80, *P* < 10^-16^). Left fPR was more correlated with face- vs tool-preferring LOTC (*t*(414) = 2.90, *P* < .005) and with face- vs tool-preferring VOTC (*t*(414) = 8.08, *P* < 10^-14^). Right fPR was more correlated with face- vs tool-preferring LOTC (*t*(414) = 3.52, *P* < .001) and with face- vs tool-preferring VOTC (*t*(414) = 8.08, *P* < 10^-14^). Left tPR was more correlated with tool- vs face-preferring LOTC (*t*(414) = 3.27, *P* < .001) and with tool- vs face-preferring VOTC (*t*(414) = 2.56, *P* < .01). Right tPR was more correlated with tool- vs face-preferring LOTC (*t*(414) = 2.79, *P* < .005), while a significant difference was not observed for VOTC (*t*(414) = 1.57, *P* = .059). Overall, these results demonstrate that category-sensitive regions with TP and PR have stronger functional connectivity with areas of LOTC and VOTC that show similar category preferences.

## Discussion

The present study used a large-scale fMRI dataset to characterize the functional organization of the anterior temporal lobe (ATL) with respect to visual category preference. Using an ROI-based approach optimized for sensitivity in a region susceptible to signal dropout, we identified three distinct profiles of visual category response within the ATL: an object- or person-preferring region in temporal pole (TP), a face-preferring region of perirhinal cortex (fPR), and an object-preferring region of perirhinal cortex (tPR). These response profiles were replicated in an independent sample of equal size. Furthermore, ATL subregions showed dissociable patterns of resting-state functional connectivity with the rest of cortex. Together, these findings demonstrate that category sensitivity—a hallmark of posterior occipitotemporal cortex—extends to the apex of the cortical visual hierarchy, and that the ATL contains functionally distinct subregions varying in category preference.

Expanding upon prior reports of object preferences in perirhinal cortex (Liang et al., 2013; Litman et al., 2009), our results demonstrate that PR contains at least two distinct subregions that vary in their response across visual categories and functional connectivity with posterior occipitotemporal regions. One subregion responds preferentially to faces, consistent with prior reports in humans and macaques for a face area within PR (Deen et al., 2024; Landi & Freiwald, 2017). These results provide converging evidence that the previously described ATL face area (Axelrod & Yovel, 2013; Lafer-Sousa et al., 2016; Rajimehr et al., 2009; Tsao et al., 2008) can be localized to PR (Collins & Olson, 2014; Deen et al., 2024; O’Neil et al., 2014). The response profile of face-preferring PR closely resembled that of well-characterized face-preferring regions of LOTC and VOTC, showing responses across categories ordered faces > bodies > tools > scenes. Together with its preferential functional connectivity to occipitotemporal face regions, these results support the view that fPR constitutes an extension of the ventral stream face processing hierarchy into the ATL (Collins & Olson, 2014; Duchaine & Yovel, 2015; Landi & Freiwald, 2017).

In addition to this face response, we identify a functionally distinct region of PR (tPR) with strong responses to tools and bodies over scenes and a weaker preference for faces over scenes. This result indicates that, like on the fusiform gyrus and in LOTC, processing of faces and non-face objects remains separated in anterior portions of the ventral stream. While responses to objects and tools in LOTC and VOTC have been extensively characterized (Bracci et al., 2012; Chao et al., 1999; Cichy et al., 2011; Grill-Spector et al., 1999; Grill-Spector et al., 1998; Haushofer et al., 2008; Kourtzi & Kanwisher, 2001), perirhinal responses to objects have not typically been reported despite the area’s established role in object processing from neuropsychological work. Like fPR for faces, tPR may constitute an anterior extension of the ventral stream object processing hierarchy, and is therefore a promising target for the study of high-level object representations.

In contrast to the well-characterized spatial organization of category-preferring regions in posterior occipitotemporal cortex, we found that face- and object-preferring subregions of PR did not show a consistent, large-scale spatial relationship. In the current analysis, face- and tool-preferring areas in LOTC and VOTC showed substantial and systematic differences in spatial position consistent with established topographic relationships (Chao et al., 1999). However, no clear spatial relationships were observed between fPR and tPR. One potential explanation for this finding is that category preferences within PR are organized in a patchy or "salt-and-pepper" manner, in which populations of neurons with different category preferences are interleaved at a fine spatial scale (Fujita et al., 1992). Such an organization might facilitate PR’s role in associating multiple objects (Murray & Richmond, 2001), which could be better served by distributed, overlapping representations than by strict spatial segregation. This organization could also help explain why category preferences in PR have been missed in many prior studies utilizing whole-brain analyses. Alternatively, it is possible that reliable spatial relationships between face- and tool-preferring subregions of PR were not observed due to lower signal quality in this region, or that their spatial relationship varies across individuals in ways that cannot be captured by group-level statistics. Future work using higher-resolution imaging approaches, methods optimized to boost ATL signal quality, or intracranial recordings will be necessary to address this question.

The response profile of TP was notably different from both PR subregions. Rather than showing responses clearly above baseline to any particular object category, TP was characterized primarily by a reduced response to scenes relative to other categories, and in the right hemisphere, a preference for animate (faces and bodies) over inanimate objects (tools). This pattern is consistent with neuropsychological findings implicating TP in person and object recognition (Damasio et al., 1990; Gentileschi et al., 2001; Gorno-Tempini et al., 2004; Snowden et al., 2011). The right-hemisphere dominance for face and body responses in TP aligns with a substantial literature demonstrating right-lateralization in occipitotemporal cortex (Engell & McCarthy, 2013; Kanwisher et al., 1997; Wang et al., 2020). This result is also consistent with prior neuroimaging and neuropsychological evidence on TP function, which have identified a hemispheric dissociation in which right TP is more involved in face processing and social cognition, while left TP is more involved in generic object processing and semantic knowledge (Damasio et al., 1990; Olson et al., 2007; Rice et al., 2018; Rice et al., 2015). Notably, the TP response profile was qualitatively similar regardless of which contrast was used to define ROIs, indicating spatial homogeneity of functional responses within TP that contrasts with PR and posterior occipitotemporal areas. Consistent with prior work, TP showed functional connectivity with transmodal regions of high-level association cortex, including the default mode network (Deen & Freiwald, 2025; Deen et al., 2024; Girn et al., 2024).

In the current study, we did not report a face-specific response in TP, as we have observed previously (Deen et al., 2024). Several factors may account for this discrepancy. First, Deen et al. (2024) found that TP responds most strongly to images of personally familiar faces, with a weaker response to unfamiliar faces over other categories. The current study did not include familiar faces, which may have elicited a stronger TP response. Second, Deen et al. (2024) used data acquisition and analysis methods optimized to counteract signal dropout in the ATL, which may have increased sensitivity to identify face-specific TP subregions. Further work with optimized methods will be needed to replicate and characterize face responses within TP.

While we discuss the current results in terms of object categories like faces and tools that made up the stimulus set, we emphasize that this is intended as an empirical description of visual response profiles, and not as a strong interpretation of the function of regions described. An ongoing debate in visual cognitive neuroscience concerns whether category-sensitive responses in the ventral stream are better explained as preferences for specific categories per se (Conway, 2018; Grill-Spector & Malach, 2004; Kanwisher, 2010; Op de Beeck et al., 2008; Peelen & Downing, 2017; Yargholi & Op de Beeck, 2023), or related to continuous dimensions that correlate with category membership (Bao et al., 2020; Contier et al., 2024; Huth et al., 2012; Konkle & Caramazza, 2013; Ritchie et al., 2025). Because the current results come from a blocked-design experiment with responses to only four categories of object, they cannot distinguish between these two interpretations. Future work exploring responses to a wide array of images will be necessary to determine whether categorical or dimensional models better capture visual responses in the ATL regions identified here.

Overall, the present study provides large-scale, replication-supported evidence that the ATL contains three functionally distinct subregions characterized by differing visual category preferences and dissociable patterns of functional connectivity. These findings demonstrate that the organizational principle of category sensitivity, long recognized as a hallmark of posterior occipitotemporal cortex, extends to the apex of the cortical visual hierarchy. As progress is made in optimized methods for imaging the ATL, further study of the regions identified here may elucidate mechanisms supporting the later stages of object processing in the brain.

## Data and code availability

Data from the young adult Human Connectome Project and are available at https://www.humanconnectome.org/study/hcp-young-adult/document/1200-subjects-data-release. Analysis code is available at https://github.com/bmdeen/fmriPermPipe (generic analysis tools) and https://github.com/SocialMemoryLab/HCP_ATL (project-specific wrapper scripts).

## Acknowledgements

This project was supported by a grant from the Louisiana Board of Regents (LEQSF[2024-27]-RD-A-25 to B.D.). Data were provided by the Human Connectome Project, WU-Minn Consortium (Principal Investigators: David Van Essen and Kamil Ugurbil; 1U54MH091657) funded by the 16 NIH Institutes and Centers that support the NIH Blueprint for Neuroscience Research, and by the McDonnell Center for Systems Neuroscience at Washington University. We thank Anisah Sahibul and Adya Agarwal for their assistance with data curation and lab operations.

## Author Contributions

B.D. conceptualized the projection, provided supervision, and developed analysis software. C.S. and B.D. analyzed the data and wrote the paper.

